# Methods for detection of cardiac glycogen-autophagy

**DOI:** 10.1101/2024.08.11.607511

**Authors:** Parisa Koutsifeli, Lorna J. Daniels, Joshua Neale, Sarah Fong, Upasna Varma, Marco Annandale, Xun Li, Yohanes Nursalim, James R. Bell, Kate L. Weeks, Aleksandr Stotland, David J. Taylor, Roberta A. Gottlieb, Lea M.D. Delbridge, Kimberley M. Mellor

## Abstract

Glycogen-autophagy (‘glycophagy’) is a selective autophagy process involved in delivering glycogen to the lysosome for bulk degradation. Glycophagy protein intermediaries include STBD1 as a glycogen tagging receptor, delivering the glycogen cargo into the forming phagosome by partnering with the Atg8 homolog, GABARAPL1. Glycophagy is emerging as a key process of energy metabolism and development of reliable tools for assessment of glycophagy activity is an important priority. Here we show that antibodies raised against the N-terminus of the GABARAPL1 protein (but not the full-length protein) detected a specific endogenous GABARAPL1 immunoblot band at 18kDa. A stable GFP-GABARAPL1 cardiac cell line was used to quantify GABARAPL1 lysosomal flux via measurement of GFP puncta in response to lysosomal inhibition with bafilomycin. Endogenous glycophagy flux was quantified in primary rat ventricular myocytes by the extent of glycogen accumulation with bafilomycin combined with chloroquine treatment (no effect observed with bafilomycin or chloroquine alone). In wild-type isolated mouse hearts, bafilomycin alone and bafilomycin combined with chloroquine (but not chloroquine alone) elicited a significant increase in glycogen content signifying basal glycophagy flux. Collectively, these methodologies provide a comprehensive toolbox for tracking cardiac glycophagy activity to advance research into the role of glycophagy in health and disease.

## Introduction

Autophagy is a ubiquitous cellular bulk degradation process that utilizes the acidic environment of the lysosome to break down damaged proteins, organelles, and macromolecules. Cellular cargo that is destined for lysosomal breakdown is tagged by autophagy receptor proteins and sequestered into double-membrane vesicles (autophagosomes) via binding to autophagy-related protein 8 (Atg8) family proteins in the phagosomal membrane.^1,2^ The autophagosome fuses with a lysosome via a coordinated process involving syntaxin, synaptobrevin and Rab family proteins. Measurement of Atg8 family proteins (in particular, LC3B) has been employed to monitor autophagy activity in mammalian cells and tissues.^3^

It is increasingly recognised that there are selective autophagy subtypes, each with specialized protein machinery to recycle specific cargo. These subtypes include macrophagy (protein macromolecules), mitophagy (mitochondria), ERphagy (endoplasmic reticulum (ER)), lipophagy (lipids), xenophagy (bacteria/viruses/pathogens) and glycophagy (glycogen).^4^ The concept of lysosomal breakdown of glycogen has been known for some time, particularly in the context of glycogen storage diseases. More recently, the specific glycophagy protein intermediaries have been identified and an understanding of the mechanism of glycogen tagging and Atg8 glycophagosome processing has been advanced in the literature.^5–8^ Glycophagy has been proposed to involve tagging of glycogen by starch-binding domain 1 (STBD1), a protein which contains both a carbohydrate binding complex (CBM20) and several Atg8 binding sites.^6,9^ STBD1 anchors to γ-aminobutyric acid receptor-associated protein-like 1 (GABARAPL1, an Atg8 homolog) in the forming phagosome membrane.^7^ STBD1 colocalization with GABARAPL1 has been demonstrated, but whether STBD1 is present in the lysosome is unclear. When the phagosome cargo is enclosed the glycophagosome fuses with a lysosome containing acid α-glucosidase (GAA) which hydrolyses the glycosidic bonds to release glucose monomers.^5^ We have previously demonstrated that glycophagy is operational in the adult heart, and is responsive to metabolic stress.^10,11^ Glycophagy is emerging as an important process of cellular energy metabolism in the heart, and early evidence suggests that it may play a role in pathological settings of cardiac metabolic stress.^12^ Thus, development and validation of methods for assessment of glycophagy in heart tissue and cardiomyocytes is an important priority.

Given that autophagy in general is a dynamic process, a major challenge for the autophagy field is experimental reliance on ‘snapshot’ measures of autophagy machinery proteins to infer regulatory response. This approach can produce contradictory findings as similar Atg8 protein levels could be measured where there is both high phagic flux and low phagic flux, because the outer-membrane Atg8 proteins are cycled into and out of phagosomes during the phago-lysosome formation cycle. Where immobilized phagosome Atg8 entrapment occurs (i.e. the Atg8 is bound to a forming phagosome which does not progress through full formation or to cargo release stage) the total protein level may not change, but autophagic flux could be decreased. Thus, assessment of autophagic activity/flux may provide a more informative indicator of autophagosome functionality.^3,13^ For macro-autophagy, immunoblot detection of the lipidated autophagosome-bound form of LC3B has provided a relatively easy approach to obtain information about pathway activity. The ratio of lipidated LC3BII to non-lipidated (cytosolic) LC3BI (LC3BII:I ratio) is commonly reported in the literature.^3^ Whether this approach can be applied to other Atg8 family proteins has not been validated. Additionally, lysosomal inhibitors have been employed as useful tools to assess autophagy flux. By blocking the downstream event of phagosome-lysosome fusion, the extent of accumulation of autophagosome protein intermediaries (e.g. LC3BII), can provide a measure of autophagy throughput/flux.

Identification of selective autophagy subtypes has been possible through the development of methodologies specific to the unique molecular signature of each subtype. The high sequence similarity of Atg8 family proteins presents a challenge in the development of antibodies which can selectively detect distinct Atg8 proteins involved in selective autophagy. For glycophagy, measurement of the Atg8 homolog, GABARAPL1, may be informative to monitor changes in activity, given its preference for partnering with the glycophagy-tagging protein, STBD1.^14^ The GABARAP Atg8 subfamily consists of GABARAP, GABARAPL1 and GABARAPL2, with relatively high sequence similarity. A previous study has highlighted the lack of specificity in some GABARAPL1 antibodies using recombinant proteins in cell lines (HEK293 cells) and mouse brain tissue.^15^ Specificity of GABARAPL1 antibodies for detection of endogenous GABARAPL1 in cardiac tissues has not been previously investigated. Glycophagy measurement has the distinct advantage that the cargo (glycogen) can be directly visualized (microscopically) and quantified by assay. The extent of glycogen accumulation in response to lysosomal inhibition can thus provide a useful tool for monitoring glycophagy flux.

The goal of this study was to validate methodological tools for monitoring glycophagy in heart tissues and cardiomyocytes. A novel GABARAPL1 knockout mouse model provided a robust negative control for validation of GABARAPL1 antibodies. Methods for assessment of glycophagy flux were evaluated in a genetically-modified cardiomyocyte cell line, primary cardiomyocytes and *ex vivo* perfused mouse hearts. This study addresses the methodologic challenges of detecting & quantifying glycophagy in general, highlights the difficulties of working with the GABARAPL1 Atg8 homologue compared with the LC3B Atg8, and offers some effective approaches for investigators working in cardiac fields.

## Results

### High sequence similarity between GABARAP Atg8 subfamily proteins compromises GABARAPL1 antibody specificity

GABARAPL1 shares high sequence similarity with the other 2 members of the GABARAP Atg8 subfamily, GABARAP and GABARAPL2 (86% and 61% similarity respectively, Fig. 1A, B). The amino acid differences between GABARAPL1 and GABARAPL2 are distributed throughout the 117 amino acid sequence (Fig. 1A). In contrast, amino acid differences between GABARAPL1 and GABARAP are more localized in the N’ terminus of the proteins (Fig. 1A). Three commercially available antibodies were tested for GABARAPL1 specificity: #1 Proteintech (11010-1-AP), a polyclonal antibody raised against the whole protein sequence; #2 Abcam (ab86497), a polyclonal antibody raised against N-terminal amino acids 1-50; #3 Cell Signaling (26632), a monoclonal antibody raised against unspecified N-terminal amino acids (Fig. 1C). To generate a negative control to use in validation of the commercially available GABARAPL1 antibodies, a Crispr/Cas9-induced global Gabarapl1 homozygous gene deletion mouse model was deployed. Guide RNAs (gRNAs) targeted sequences within the first intron and the exo-genic region post the fourth exon for Cas9-mediated excision (Fig. 1D). Thus 3 of 4 exons of the Gabarapl1 gene were deleted (6,052bp, genome annotation release GRCm38.p4). Deletion of the Gabarapl1 gene was verified via PCR (Fig. 1E).

**Figure 1.**
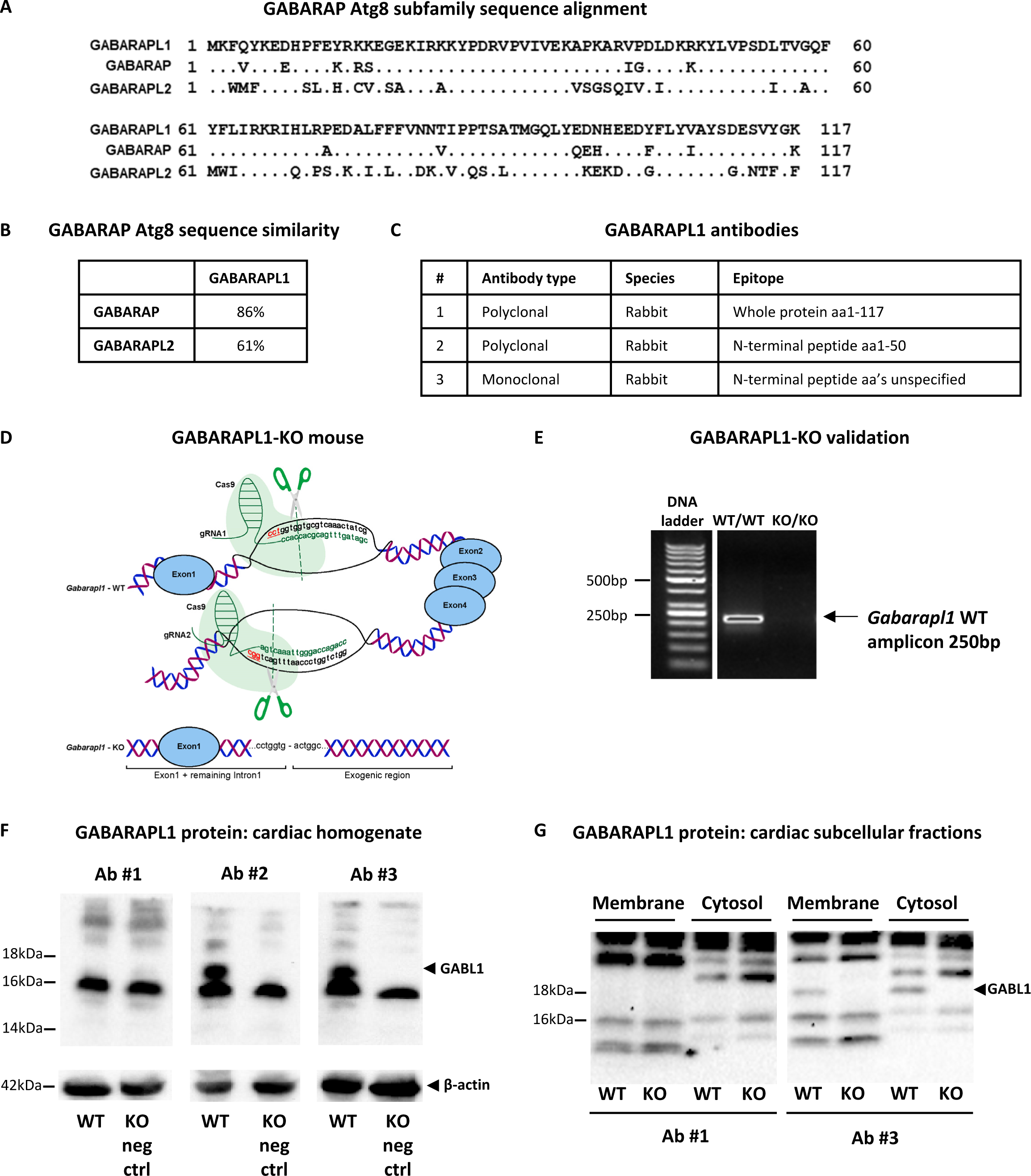
High sequence similarity between GABARAP Atg8 proteins compromises GABARAPL1 antibody specificity. A. Protein sequence alignment of human GABARAP Atg8 subfamily members. **B.** Percentage similarity between protein sequences of GABARAP, GABRAPL2 and GABARAPL1. **C.** Characteristics of three commercially available GABARAPL1 antibodies (#1 Proteintech (11010-1-AP), #2 Abcam (ab86497), #3 Cell signaling (26632)). **D.** Schematic of Crispr/Cas9 genome editing design targeting Gabarapl1 for gene deletion in mice (green: homologous part of gRNA sequences, red: proto-spacer adjacent motifs). **E.** Validation of Gabarapl1 knockout using DNA electrophoresis (tail sample). **F.** Immunoblot of total cardiac homogenate shows that Ab#2 and Ab#3, but not Ab#1, detect a specific GABARAPL1 band at 18kDa, validated using the GABARAPL1-KO mouse heart as a negative control (note absence of 18kDa GABARAPL1 (GABL1) band in KO samples for Ab#2 and Ab#3, but not Ab#1). **G.** Immunoblot of crude membrane (containing autophagosome membrane structures) and cytosol fractions from cardiac homogenate samples comparing Ab#1 and Ab#3. A specific 18kDa GABARAPL1 band is detected in membrane and cytosol fractions using Ab#3 (validated by KO sample). Ab#1 does not detect the 18kDa GABARAPL1 band in either fraction.

Total cardiac homogenate from wild type or homozygous GABARAPL1-KO mice was used to test the three commercially available GABARAPL1 antibodies using immunoblotting. An 18kDa GABARAPL1 band was detected with antibody #2 and #3, but not with antibody #1 (Fig. 1E). The absence of this 18kDa band in the GABARAPL1-KO samples confirmed that it was specific for GABARAPL1. A prominent non-specific 16kDa band was detected with all antibodies (Fig. 1E). Adequate separation of proteins in the electrophoresis gel was essential to distinguish the specific 18kDa band from the non-specific 16kDa band. These findings suggest that antibodies raised against the N-terminus of the GABARAPL1 protein (Ab#2 and 3) exhibit higher specificity for GABARAPL1 than antibodies raised against the whole protein (Ab#1).

Quantification of the lipidated ‘active form’ of the Atg8 homolog LC3B (phagosome-localized LC3BII) is commonly used as an indicator of macro-autophagy activity. Lipidation induces an apparent 2kDa shift in LC3B band size on immunoblot, thus the ratio of lipidated (phagosome-bound) LC3BII to cytosolic LC3BI can be calculated. It could be speculated that incorporation of GABARAPL1 into the phagosome membrane via lipidation would produce a similar separation in immunoblot bands. However in the total cardiac homogenate samples tested in Fig. 1E, only one specific GABARAPL1 band could be detected. Potentially, smaller kDa GABARAPL1 bands were obscured by the prominent non-specific 16kDa band.

To localize intracellular lipidated ‘active’ GABARAPL1 protein in membrane-enriched samples, total cardiac homogenate was fractionated to obtain the triton-soluble ‘crude membrane’ fraction, and the cytosolic fraction. Similar to the total cardiac homogenate blot, antibody #1 did not detect any specific GABARAPL1 bands (all bands detected were also present in the KO negative control, Fig. 1G). Antibody #3 detected a specific 18kDa GABARAPL1 band in both the crude membrane and cytosolic fractions (Fig. 1G). No other specific bands were detected, suggesting that it is not feasible to separate the lipidated vs non-lipidated forms of GABARAPL1 in these cardiac samples. Although fractionation provides a clearer view of the 18kDa band of interest, the lipidated form of GABARAPL1 may remain obscured by non-specific bands. Alternatively, it is possible that the 18kDa GABARAPL1 band corresponds to both lipidated and non-lipidated GABARAPL1 forms as it was detected in both membrane and cytosol fractions (as lipidation effects on gel migration can be complex).

### Quantification of cardiac glycophagy flux in vitro and ex vivo

Macro-autophagy flux is commonly measured by the extent of accumulation of lipidated LC3B (LC3BII) protein in response to lysosomal blockade. Figure 2A illustrates the site of action for lysosomal inhibitors chloroquine and bafilomycin. Lysosomal uptake of chloroquine (a weak base) raises the pH of the lysosome and prevents autophagosome-lysosome fusion.^16^ Bafilomycin inhibits the V-ATPase, thus preventing lysosomal acidification by the proton pump.^3^

**Figure 2.**
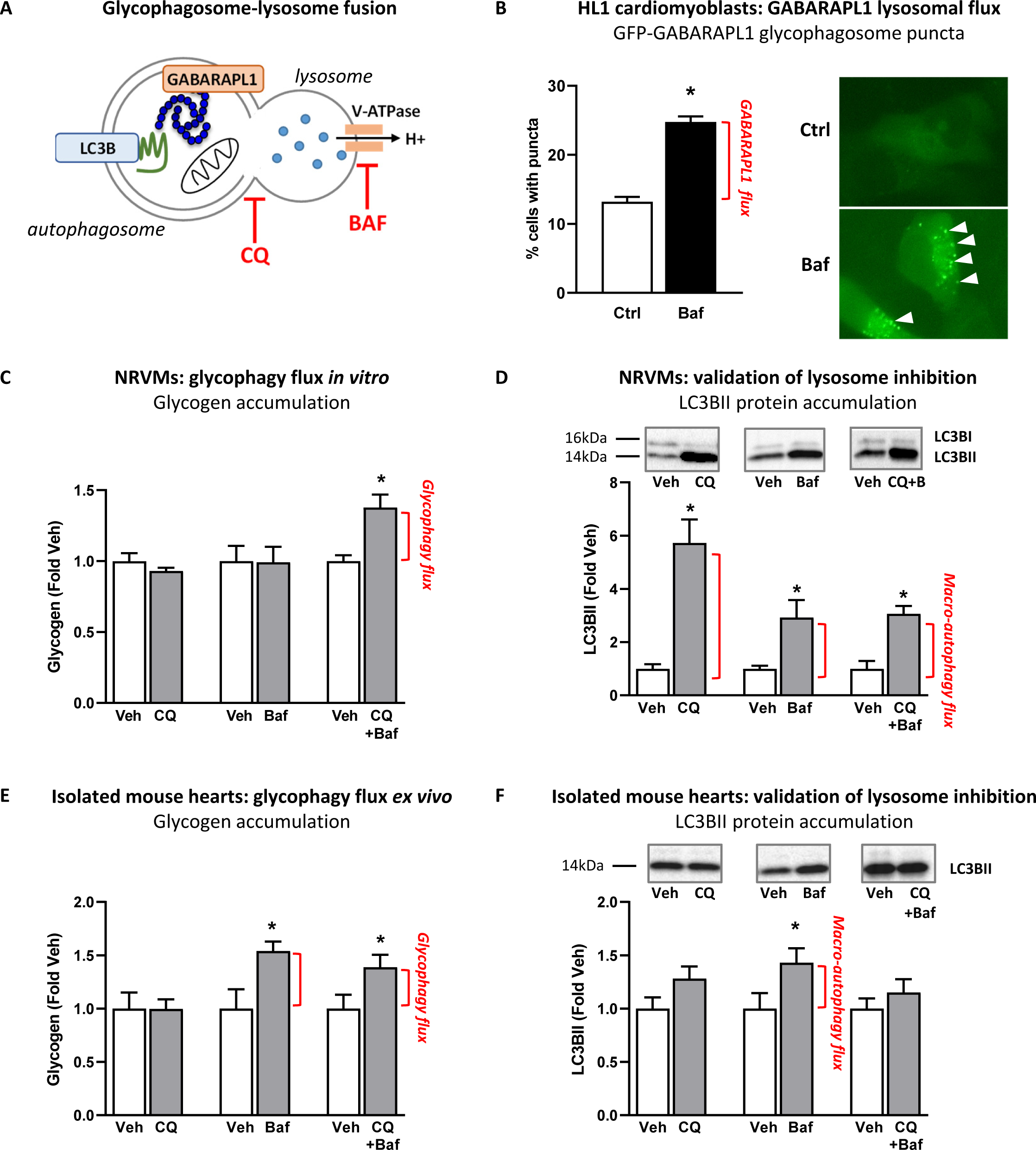
Quantification of cardiac glycophagy flux *in vitro* and *ex vivo*. A. Autophagosome-lysosome fusion schematic illustrating the mechanism of action of lysosomal inhibitors chloroquine (CQ) and bafilomycin (Baf). **B.** Glycophagosome membrane-bound GABARAPL1 puncta visualized using GFP-Gabarapl1 in mouse atria-derived HL1 cardiomyocytes. HL1 cells were transduced with GFP-Gabarapl1 and imaged before (control, Ctrl) and after 2 hours lysosomal inhibition with 100 nM Baf. Lysosomal Gabarapl1 flux is determined by the extent of accumulation of GFP-Gabarapl1 puncta in response to lysosomal inhibition with Baf. Quantification of GFP epifluorescent micrographs presented as the number of puncta-containing cells as a percentage of total cells/60x image, puncta are indicated by white arrows (9-13 images/well, n=3 independent wells). **C.** Glycophagy flux is measured in neonatal rat ventricular myocytes (NRVMs) following 2h treatment with 50 µm CQ, 100 nM Baf or 50 µm CQ + 100 nM Baf. Flux is determined by the extent of glycogen accumulation in response to lysosomal inhibition (n=5-11 independent wells per group). **D.** Immunoblot showing accumulation of macro-autophagy marker LC3BII confirms lysosomal inhibition by CQ, Baf and CQ + Baf treated NRVMs (n=6-9 independent wells per group). **E.** Glycophagy flux is measured in *ex vivo* isolated perfused mouse hearts following 30min treatment with 20 µm CQ, 1 µm Baf or 20 µm CQ + 1 µm Baf. Flux is determined by the extent of glycogen accumulation in response to lysosomal inhibition (n=5-10 mice per group). **F.** Immunoblot showing accumulation of macro-autophagy marker LC3BII confirms lysosomal inhibition by Baf, but not CQ or CQ + Baf in perfused mouse hearts (n=5-10 mice per group). Data presented as mean ± SEM, *p<0.05.

Given our demonstration that it is not feasible to monitor cardiac glycophagy activity by measurement of lipidated ‘active’ GABARAPL1 protein levels (see Fig. 1E-F), we sought to establish a novel *in vitro* approach to track glycophagosome GABARAPL1 traffic using a murine atria-derived cardiomyocyte cell line (HL1), stably expressing GFP-tagged GABARAPL1. Live cell epifluorescence imaging was used to identify GFP-GABARAPL1 puncta, representing glycophagosome-bound GABARAPL1. In response to bafilomycin, the percentage of cells with punctate GFP-GABARAPL1 distribution was significantly increased (control: ∼13% vs bafilomycin: ∼25%, Fig. 2B). The extent of increase in cells with glycophagosome-bound GABARAPL1 in response to bafilomycin-induced lysosomal blockade provides an indicator of lysosomal GABARAPL1 flux in this *in vitro* cell line setting.

To develop a method to monitor endogenous glycophagy flux in primary cardiomyocytes, neonatal rat ventricular myocytes (NRVMs) were isolated and cultured in chloroquine or bafilomycin alone, or combined bafilomycin and chloroquine for 2 hours prior to cell lysis and glycogen quantification. Surprisingly, cardiomyocyte glycogen content was unchanged in response to chloroquine or bafilomycin alone (Fig. 2C, Table S1), despite evidence of successful lysosomal blockade demonstrated by a dramatic increase in LC3BII protein (5.7-fold increase in LC3BII with chloroquine, 2.9-fold with bafilomycin, Fig. 2D). In contrast, combined bafilomycin and chloroquine treatment induced a 38% increase in glycogen content (Fig. 2C, Table S1). As expected, LC3BII levels were significantly elevated in this setting (Fig. 2D). Taken together, these findings suggest that detection of endogenous cardiomyocyte glycophagy flux requires a ‘double-hit’ of lysosomal blockade.

Validated tools for monitoring glycophagy flux in an *in vivo* setting are important for advancing understanding of the role of glycophagy in disease states. *In vivo* administration of lysosomal inhibitors has highly variable cardiac outcomes, especially in mice. Therefore, we tested a method to measure glycophagy flux in isolated mouse hearts perfused in Langendorff mode. After an initial 20 min stabilization period, hearts were perfused with bafilomycin or chloroquine (vs. vehicle control: DMSO or saline respectively), or bafilomycin and chloroquine combined, for 30 min. A significant increase in cardiac glycogen content was evident in response to bafilomycin alone and bafilomycin combined with chloroquine, but not chloroquine alone (Fig. 2E). Lysosomal inhibition with bafilomycin significantly increased LC3BII protein content with bafilomycin, but surprisingly LC3BII accumulation did not reach significance with chloroquine or with chloroquine and bafilomycin combined (Fig. 2F). Glycogen synthase and glycogen phosphorylase were unaffected by lysosomal inhibition (Fig. S1). Collectively, these findings suggest that bafilomycin is more effective than chloroquine as a tool for measuring glycophagy flux in isolated perfused mouse hearts.

## Discussion

This report provides the first comprehensive evaluation of tools for assessing cardiac glycophagy, in several experimental settings - *in vitro, ex vivo* and *in vivo*. Three antibodies targeting the glycophagosome protein, GABARAPL1, were tested using our novel *in vivo* global GABARAPL1-KO mouse model as a robust negative control. Two of the three antibodies were validated for specific detection of endogenous GABARAPL1 in murine cardiac samples. Using a subcellular fractionation approach, we determined that the lipidated (glycophagosome-bound) form of GABARAPL1 could not be distinguished from the non-lipidated form using immunoblotting, at least in a cardiac setting. Thus, alternative methods for monitoring glycophagy activity using pharmacological lysosomal inhibition were pursued. A stable cardiac cell line with GFP-tagged GABARAPL1 was validated as an *in vitro* tool for quantifying GABARAPL1 lysosomal flux via measurement of GFP puncta in response to lysosomal inhibition with bafilomycin. An *in vitro* method for monitoring endogenous glycophagy flux was validated in neonatal rat ventricular myocytes involving measurement of glycogen accumulation in response to lysosomal inhibition with combined bafilomycin and chloroquine (but not with bafilomycin or chloroquine alone). And finally, an *ex vivo* method for monitoring glycophagy flux was validated in isolated mouse hearts perfused with bafilomycin or bafilomycin combined with chloroquine (but not chloroquine alone) by measurement of the extent of glycogen accumulation in response to lysosomal inhibition. These studies characterize the utility of selected tools for monitoring glycophagy in response to interventions performed *in vitro* or *in vivo*. Conventional methods for detection of macro-autophagy flux are not necessarily valid for selective autophagy subtypes – specifically glycophagy, as we show here. The differential efficacy of the lysosomal inhibitors bafilomycin and chloroquine to modulate glycophagy activity in various experimental settings highlights the importance of validating experimental tools available for specific investigative contexts.

The generation of specific GABARAPL1 antibodies is complicated by the high sequence similarity between GABARAP family proteins: in particular the close alignment of GABARAPL1 and GABARAP (86% similarity). In the present study, we demonstrate that antibodies raised against the N-terminus of the GABARAPL1 protein (Ab#2 Abcam #ab86497 and Ab#3 Cell Signaling #26632) detect a specific endogenous GABARAPL1 immunoblot band at 18kDa. The antibody raised against the full length of the GABARAPL1 protein (Ab#3 ProteinTech 11010-1-AP) did not detect any specific bands and is therefore not validated for use in this cardiac setting. Previous studies have shown that this antibody can distinguish between GABARAPL1 and GABARAP when comparing HEK293 cells transfected with either GFP-tagged recombinant GABARAPL1 or GABARAP.^15^ The same study also showed that the ProteinTech 11010-1-AP GABARAPL1 antibody specifically stained GABARAPL1 in rat brain sections, using a pre-incubation antibody blocking approach as a negative control.^15^ Thus it is feasible that in experimental settings where the migration of GABARAPL1 protein is shifted in the electrophoresis gel (i.e. via addition of a GFP tag), or in tissues where the relative expression of GABARAPL1 is high (i.e. rat brain), this antibody may specifically detect GABARAPL1. In contrast, in an endogenous cardiac setting where prominent non-specific proteins migrate to locations in the SDS-PAGE gel which are very similar to GABARAPL1, usage of antibodies raised against the N-terminus of the GABARAPL1 protein are more suitable.

Given that autophagy is a dynamic process, snapshot measures of autophagy proteins may not necessarily capture the alterations in autophagic activity in response to an intervention. Thus, the ratio of the lipidated (autophagosome-bound, LC3BII) to non-lipidated (cytosol, LC3BI) form of the Atg8 protein LC3B has been frequently employed as a marker of autophagic activity. Lipidation of LC3B induces a 2 kDa shift in protein migration in an SDS-PAGE gel and appears as a distinct immunoblot band below non-lipidated LC3BI. Some studies have shown a similar band shift for GABARAPL1, but have either not provided validation of the specificity of the endogenous bands detected,^17^ or have used cells with genetically-modified GABARAPL1 inducing a molecular weight shift in the band of interest (i.e. with the addition of a FLAG-tag).^18^ In the present study, only one specific GABARAPL1 band was detected (18kDa) in total homogenate, crude membrane and cytosolic samples from mouse hearts. It is not clear whether this band represents the lipidated or non-lipidated form of the protein. In a previous study using HEK293 cells transduced with FLAG-GABARAPL1, the lipidated form of GABARAPL1 was not detected in untreated control cells, and appeared in response to lysosomal inhibition (NH_4_Cl or bafilomycin) and/or autophagy inducers (rapamycin or starvation medium).^18^ Overexpression of Atg4B, a cysteine protease which delipidates GABARAP family Atg8 proteins,^19^ decreased intensity of the band corresponding to lipidated FLAG-GABARAPL1.^18^ These findings suggest that GABARAPL1 is lipidated in a similar manner to LC3B, which may induce a shift in the migration of the protein in an SDS-PAGE gel. In the mouse heart samples investigated in the present study, several non-specific bands are detected in a similar gel location to that expected for a 2 kDa lipidation shift. Thus the use of a GABARAPL1-II:I lipidation ratio as a tool for monitoring endogenous cardiac glycophagy activity is not feasible in this setting, although it might be possible to quantify the amount of GABARAPL1 in cytosol vs. membrane fractions normalized to cell equivalents.

Lysosomal inhibitors have been employed as a tool to measure autophagy flux *in vitro* and *in vivo*.^3,13^ Quantification of the extent of accumulation of LC3BII (lipidated, autophagosome-bound form) in response to acute lysosomal inhibition provides a measure of LC3BII lysosomal degradation, indicative of macro-autophagy throughput. Given that the lipidated form of GABARAPL1 is not selectively detectable by immunoblot in cardiac samples, alternative endpoint measures were pursued in the present study to develop a method to quantify glycophagy flux in cultured cardiomyocytes *in vitro*, and mouse hearts *ex vivo*. A novel GFP-tagged GABARAPL1 stable cardiomyocyte cell line (HL1 murine atria-derived cells) was established to visualize the capture of GABARAPL1 into glycophagosomes (punctate appearance) in response to bafilomycin-induced lysosomal inhibition.

To monitor endogenous glycophagy flux levels, a primary cardiomyocyte culture setting was employed, using cellular glycogen content as an endpoint. The extent of glycogen accumulation in response to lysosomal inhibition provides a measure of glycophagy flux. Interestingly, neither chloroquine or bafilomycin alone were sufficient to detect basal glycophagy flux in quiescent primary cells, despite eliciting a robust increase in LC3BII. These findings suggest that tools applied for detection of macro-autophagy flux are not necessarily transferrable to quantification of glycophagy flux. The use of glycogen as an endpoint measure relating to phagosome ‘cargo’ processing (compared with Atg8 lipidation indicative of phagosomal localization) appears to require a ‘double-hit’ of bafilomycin and chloroquine. Given that glycophagy is involved in liberating glucose fuel for cellular glycolytic metabolism,^5^ baseline glycophagy flux may be relatively low in a quiescent cell culture setting of minimal metabolic demand. In contrast, in the *ex vivo* context of a rapidly paced heart perfusion, bafilomycin alone was sufficient to detect glycophagy flux, as determined by the extent of glycogen accumulation following 30 minutes lysosomal inhibition.

Chloroquine alone did not elicit a glycogen or significant LC3BII response, suggesting that lysosomal inhibition was not successfully induced in this setting. Previous studies have also shown that LC3B is unchanged in response to chloroquine perfusion in isolated rat hearts.^20^ In the present study, chloroquine perfusion decreased myocardial coronary flow rate (data not shown) which may have suppressed metabolic demand and obviated differences in glycogen and LC3B induced by lysosomal inhibition. This observation is consistent with a previous report that chloroquine decreased left ventricular contractility in isolated rat hearts,^21^ and may reflect vascular-related side-effects.

Given that chloroquine is well described to elicit autophagy-independent actions,^22^ it is possible that the off-target effects of chloroquine may have influenced the glycogen response in this study. In L6 muscle cells and rat skeletal muscle tissue, it is reported that chloroquine activated Akt signalling and promoted glucose uptake and dephosphorylation (activation) of glycogen synthase.^23^ In the present study, if chloroquine-induced activation of glycogen synthase was dominant, an exacerbation of the glycogen response would be expected. Interestingly, glycogen levels were unchanged in primary cardiomyocytes and isolated hearts treated with chloroquine alone, suggesting that off-target promotion of glycogen synthesis by chloroquine was not evident in this setting.

Alternative approaches for monitoring glycophagy have been reported in the literature, each with inherent limitations:

*i) Lysosomal acid α-glucosidase (Gaa) activity assay.*^24,25^ While Gaa activity is certainly informative in relation to the end-stage of the glycophagy process, it may not necessarily track with glycophagy throughput in all settings. For example, upregulation of the rate of Gaa-mediated glycogen breakdown may be a compensatory response to a low rate of glycophagosome glycogen capture. It is not yet known where the rate-limiting steps are located in the glycophagy pathway, but in most biological processes the rate of throughput is determined at early and intermediate steps. Future studies tracking Gaa activity against glycophagy flux measures are required to validate whether findings relating to Gaa enzyme status can be extrapolated to conclusions about glycophagy throughput.
*ii) Quantification of glycophagosomes evident in electron micrographs.*^26^ Transmission electron microscopy is an excellent approach to visualize glycophagosomes, due to the intrinsic high electron density of glycogen particles which is further enhanced with lead citrate staining. But given that glycophagosomes are only discernible at very high magnification, the reliability of quantitation is limited – particularly in adult samples where glycophagosome abundance is lower than the neonatal setting. Achieving visualization and quantitation of the phagosome and the phagolysosome stages of conventional autophagy (i.e. macroprotein autophagy) remains an important challenge in the broader autophagy research field. The stochastic nature of autophagy events and the requirement for high magnification EM to identify the small phagosome structures make data sampling and quantitation inherently problematic. Regardless, an increase in glycophagosome quantity could reflect either an accumulation of glycophagosomes due to pathway impairment, or an upregulation of the process in settings of high glycophagy flux.
*iii) Co-localization of GABARAPL1 and glycogen (co-immunostaining)*^27^ *, lysosomal proteins and glycogen, or of GABARAPL1 and STBD1 (co-immunoprecipitation).*^28–30^ These data would provide a static measure of glycogen in the lysosome and not glycophagy flux per se. Increased colocalization may reflect increased or decreased flux. These approaches are reliant on the specificity of the antibodies employed. In the present study we have demonstrated that even if the GABARAPL1 immunoblot band can be validated (via use of our KO model), there are prominent non-specific bands in other molecular weight ranges. Thus it is highly likely that the available GABARAPL1 antibodies would generate off-target fluorescence in co-immunostained sections and pull down non-specific proteins in immunoprecipitation processes. Similarly, in our hands STBD1 antibodies are also unsuitable for immunoprecipitation due to detection of several prominent non-specific bands on immunoblot, albeit at a higher molecular weight range than the size of STBD1. Analysis of changes in STBD1 subcellular localization during manipulation of glycophagy would be informative, and further development of methods to achieve this is warranted.
*iv) Use of GAA inhibitors to block lysosomal glycogen degradation and estimate glycophagy flux.* GAA inhibitors have limited cell uptake, thus use as an agent to inhibit subcellular processes is limited.
*v) Measurement of lysosomal glycogen content.* Lysosomal glycogen content would be expected to be below the limit of detection of standard glycogen assays, particularly for adult mouse hearts where glycogen content is relatively low.

In conclusion, our findings confirm the specificity of two GABARAPL1 antibodies raised against the N-terminus of the GABARAPL1 protein and show that detection of lipidated vs non-lipidated GABARAPL1 is not feasible in an endogenous cardiac setting. We demonstrate three methods for monitoring cardiomyocyte glycophagy flux using a novel GFP-GABARAPL1 cardiomyocyte cell line, endogenous primary cardiomyocytes and perfused mouse hearts. We compare and contrast the effectiveness of lysosomal inhibitors, chloroquine and bafilomycin, for utility in measurement of glycophagy (vs macro-autophagy) flux. In primary cells, combined bafilomycin and chloroquine treatment are required to elicit a glycogen response, while bafilomycin perfusion alone is sufficient for glycogen accumulation in isolated mouse hearts. It is apparent that the tools required for assessment of glycophagy flux should be tailored to the investigative setting and that strategies for quantifying macro-autophagic flux may not be effective for other selective autophagic processes. Whether these methods are transferrable to non-cardiac cell types requires validation.

Collectively, the methodologies reported here extend the autophagy ‘tool-kit’ for specific tracking of glycophagy activity. We establish a ‘negative control’ process suitable for testing new GABARAPL1 immuno-reagents, and our findings offer potential to expand experimental capacity to advance research into the role and regulation of glycophagy in the heart in health and disease. It is anticipated that usage of glycogen-flux methodologies described here will assist in evaluating glycogen levels in cardiac and other cell types when expression of glycogen handling proteins such as GABARAPL1 are disrupted.

## Materials and Methods

### Animals

Animal experiments were conducted at the University of Auckland and the University of Melbourne in accordance with the *Good Practice Guide for the Use of Animals in Research* and the *Australian Code for the Care and Use of Animals for Scientific Purposes* (NHMRC, 2013). Animals were housed in the Vernon Jensen Unit (Auckland) or Biomedical Sciences Animal Facility (Melbourne). Animals were maintained under temperature controlled conditions and a 12:12 hour light:dark cycle. Food and water were provided *ad libitum*.

### Crispr-Cas9 GABARAPL1-KO mice, DNA extraction and genotyping

The founder generation of Crispr-Cas9 global GABARAPL1-KO mice was generated by the Melbourne Advanced Genome Editing Centre. Briefly, C57/Bl6J embryos were injected with Cas9 from Streptococcus pyogenes mixed with gRNAs targeting the *Gabarapl1* gene (gRNA-1: GGTGGTGCGTCAAACTATCG, gRNA-2: GGTCTGGTCCCAGATTTGAC) and were subsequently implanted into pseudo-pregnant females. Founder GABARAPL1-KO mice were identified by Sanger sequencing (Geneious software v11.0.5, Biomatters Ltd) and were crossed with wild type C57Bl/6J mice for 2 generations. Subsequent genotyping was performed via PCR amplification following DNA extraction. Briefly, DNA was extracted from tail clips following incubation with DirectPCR Lysis reagent (Viagen, #102-T) supplemented with PCR-grade Proteinase K (Sigma, #P6556), at 55⁰C. DNA was collected via NaCl (Sigma, #S6191) and isopropanol (Sigma, #I9516) precipitation. Extracted DNA was resuspended in 10mM Tris-HCl, pH 8.0 (Sigma, #10812846001) and combined with GoTaq MasterMix (Promega, #M7833) and primers for PCR amplification. Genotypes were visualised via DNA gel electrophoresis. *Gabarapl1* primers: FWD 5’-AGTGGACATCGAAGGACAG - 3’, REV 5’ – TCAGTGAGTAAAGGCTTGCC - 3’. For tissue collection, WT and KO mice were anaesthetized (sodium pentobarbitone, 60 mg/kg i.p.) and a thoracotomy was performed. Hearts were rapidly excised and snap frozen in liquid N_2_ for later molecular analysis.

### Murine atria-derived HL1 cardiomyocyte GFP-GABARAPL1 cell line

To establish a stable HL1 cardiomyocyte cell line expressing GFP-tagged GABARAPL1, non-replicative Moloney’s murine leukemia self-inactivating retroviral particles encoding GFP-*Gabarapl1* were mixed with 5 µg/mL polybrene (Sigma), added to naïve HL1 cells, and centrifuged at 1500 g, 32 ⁰C for 80 minutes.^31^ The transduced cells were selected by 10 µg/mL blasticidin S HCl (ThermoFisher) for 6 days. HL1-GFP-GABARAPL1 cells were cultured in Claycomb media (Sigma) supplemented with 2 mM L-glutamine (Life Technologies), 10 % NBCS (Thermo Fisher Scientific), 0.1 mM norepinephrine (Sigma) and antiobiotic-antimyotic reagent (Life Technologies), as previously described.^32,33^ An epifluorescence microscope (60x oil objective, Keyence Biorevo) was used to image GFP fluorescence before (basal) and after exposure to lysosomal inhibitor, bafilomycin (100 nM, 2 hour). ImageJ software (NIH) was used to evaluate GFP-GABARAPL1 puncta. Cells were categorized as either exhibiting predominantly diffuse GFP-GABARAPL1 fluorescence or numerous punctate GFP-GABARAPL1 structures (representing GABARAPL1 accumulation in autophagosomes), and the percentage of cells with GFP-GABARAPL1 punctate was calculated for each image as previously described.^34^ Gabarapl1 lysosomal degradation is determined by the extent of accumulation of GFP-Gabarapl1 puncta in response to lysosomal inhibition with Baf.

### Cultured neonatal rat ventricular myocytes

Neonatal rat ventricular myocytes (NRVMs) were isolated from p1-2 Sprague Dawley rats (equal male and female) following collagenase digestion as previously described.^11^ NRVMs were seeded in 6-well plates at density 1,187,500 cells/well. Cells were cultured in minimum essential media (MEM, Invitrogen, #61100-061) supplemented with 10% newborn calf serum (NBCS, Invitrogen, #16010-159) for 24 hours prior to culture in Dulbecco’s modified essential medium (DMEM) supplemented with 5mM D-glucose (Sigma, #G7021) for 24 hours. For assessment of endogenous glycophagy flux, cell media was supplemented with either 100 nM bafilomycin A1 (Sapphire Bioscience Pty Ltd # S1413), 50 µM chloroquine diphosphate (Sigma, #C6628) or 100 nM bafilomycin A1 and 50 µM chloroquine diphosphate for 2 hours. Cells were lysed in RIPA buffer containing protease (Complete, EDTA-free, Roche, #04693132001) and phosphatase inhibitors (PhosSTOP, Roche, #04906837001) for molecular analysis.

### *Ex vivo* mouse heart perfusions

C57Bl/6 male mice were anaesthetized (sodium pentobarbitone, 60 mg/kg i.p.) and a thoracotomy was performed. Hearts were rapidly excised and arrested in cold (4°C) Krebs-Henseleit bicarbonate buffer (in mM: 119 NaCl, 10 glucose, 22 NaHCO_3_, 4 KCl, 1.2 MgCl_2_, 1.2 KH_2_PO_4_, 0.5 EDTA, 2 Na pyruvate). The aorta was cannulated and the heart was retrogradely perfused with oxygenated (95% O_2_-5% CO_2_) Krebs-Henseleit bicarbonate buffer (37.0°C, pH 7.4, with the addition of 2.5 mM CaCl_2_) in non-recirculating Langendorff mode at a constant pressure (80 mmHg equivalent, 10 Hz pacing), as previously described.^35^ Hearts were stabilized for 20 minutes prior to 30 minutes perfusion with 1 µM bafilomycin A1 (in 0.01% dimethylsulfoxide (DMSO)), 20 µM chloroquine diphosphate (in saline), 1 µM bafilomycin A1 combined with 20 µM chloroquine diphosphate or vehicle control (0.01% DMSO or saline respectfully), and ventricles were dissected and snap frozen in liquid N_2_ for later molecular analysis.

### Glycogen content

Glycogen content was determined in a 2-step enzymatic calorimetric assay as described previously.^11^ Briefly, duplicate sample aliquots were incubated at 50 ⁰C in the presence or absence of amyloglucosidase (Roche, # 10102857001) in 1 % triton-X, 0.1M sodium acetate, pH 6.0 for 1 hour prior to centrifugation at 16,000 g, 4 ⁰C, for 2 minutes. The supernatant was incubated with glucose oxidase/peroxidase (PGO enzyme capsules, Sigma, #P7119) with 1% o-dianisdine dihydrochloride (Sigma, #D3252) for 30 minutes at room temperature. Absorbance was measured at 450nm. Background glucose content (measured in the absence of amyloglucosidase) was subtracted from the amyloglucosidase-incubated sample, and glycogen content was presented as glucose units (nmol) normalised to protein (mg, determined by Lowry assay^36^).

### Subcellular fractionation

Frozen tissue was homogenized in lysis buffer (100 mM Tris-HCl (pH 7.0), 5 mM EGTA, 4 mM 4 ⁰C). The supernatant was collected for the cytosolic fraction and the pellet was resuspended in lysis buffer with 1 % triton-X (Sigma) and centrifuged at 16,000 g (10 minutes, 4 ⁰C). The triton-soluble supernatant was collected for the crude membrane fraction.

### Immunoblot

Protein concentration was determined by Lowry assay and samples were diluted in loading buffer (50 mM Tris-HCl (pH 6.8), 2 % sodium dodecyl sulfate, 10 % glycerol, 0.1 % bromophenol blue and 2.5 % 2-mercaptoethanol). Samples were loaded onto polyacrylamide gels (Biorad precast 4-15% acrylamide gels, #4561083) with equal protein loading and separated by SDS-PAGE gel electrophoresis prior to transfer to polyvinylidene difluoride membranes (Trans-Blot® Turbo™ Transfer System, Biorad, #1704274). Membranes were subjected to blocking and antibody incubation as previously described.^37^ For GABARAPL1 blots, membranes were blocked in 5 % bovine serum albumin (BSA) in tris buffer saline containing 0.1% Tween20. Antibodies were sourced for GABARAPL1 (#1: Proteintech, #11010-1-AP; #2 Abcam, #ab86497; #3: Cell Signaling, #26632), LC3B (Cell Signaling #2775), β-Actin (Cell Signaling, #4967), phosphorylated (Ser641) glycogen synthase (ab81230, Abcam), glycogen synthase (3893, Cell Signaling), phosphorylated (Ser14) glycogen phosphorylase (gift from Dr David Stapleton), and glycogen phosphorylase (gift from Dr David Stapleton). Membranes were incubated with anti-rabbit horseradish peroxidase conjugated secondary antibody (GE Healthcare). The ECL Prime (Amersham, GE Healthcare) chemiluminescent signal was visualized with a ChemiDoc-XRS Imaging device and band intensity quantified using ImageLab software (Bio-Rad). Equal protein loading was confirmed by Coomassie staining (Coomassie Brilliant Blue R-250, Bio-Rad). For analysis of LC3BII response to lysosomal blockade (CQ/Baf), quantitation of LC3BII bands was normalized to total protein content determined by image analysis of Coomassie-stained blots.

### Statistical analysis

Data are presented as means ± S.E.M. Data were analysed by Student’s T-Test (comparison between 2 groups) or 1-way ANOVA (comparison between 3 or more groups) with Bonferroni post-hoc analyses where appropriate (Graphpad Prism Software v.8). Parametric assumptions of normal distribution (Shapiro-Wilk test) and equal variance (F-test or Brown-Forsythe test) were verified for all datasets prior to applying parametric statistical tests. A *P-*value < 0.05 was considered statistically significant.

## Disclosure statement

The authors have no conflicts of interest to declare.

## Funding

The research was supported by funding from the Health Research Council of New Zealand (19/190), Marsden Fund, New Zealand (19-UOA-268, 14-UOA-160), National Institutes of Health USA (NIH: P01HL-112730, R01-HL132075, R01-HL144509).

## Supplementary Data

**Figure S1.**
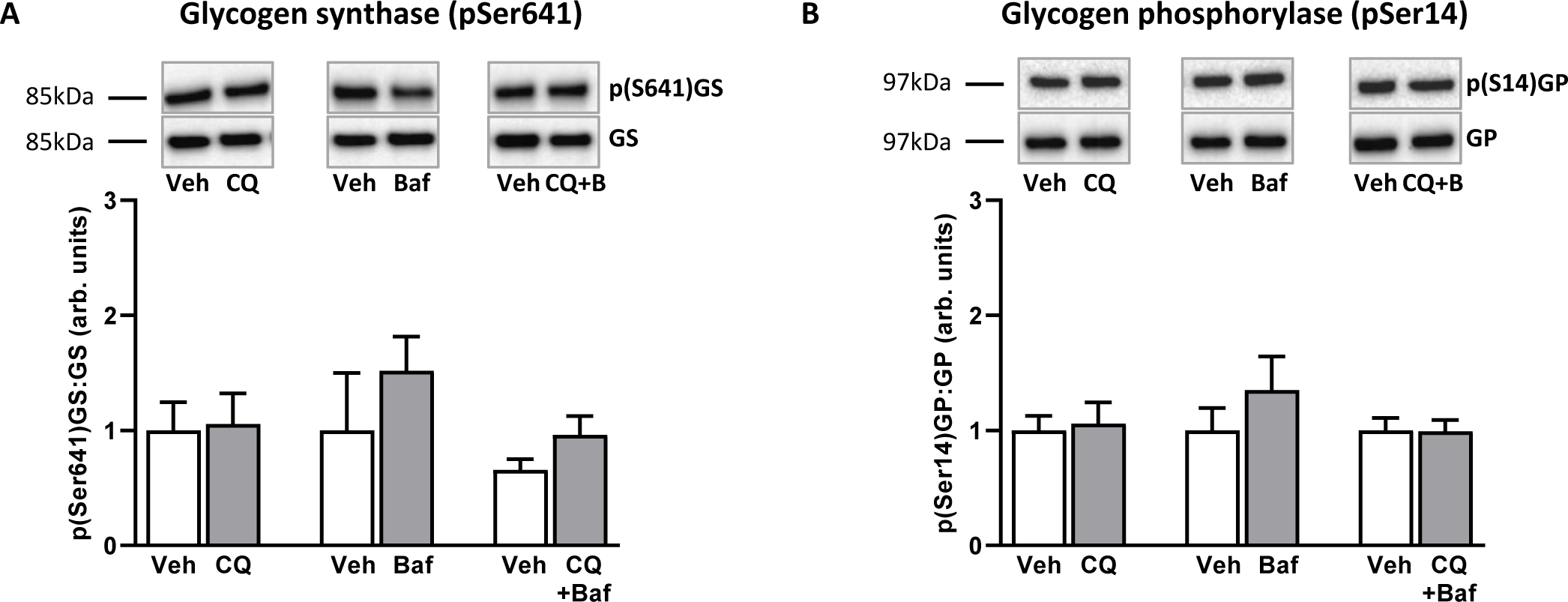
Glycogen synthase and glycogen phosphorylase in mouse hearts perfused with lysosomal inhibitors. A. Ratio of phosphorylated (Ser641) to total glycogen synthase. Note that pSer641 is an inhibitory site. **B.** Ratio of phosphorylated (Ser14) to total glycogen phosphorylase. (n=4-11 mice per group). Data presented as mean ± SEM, *p<0.05.

**Table S1.** Glycogen values in NRVM cultures.

| Group | Glycogen (nmol/mg) |
| --- | --- |
| Veh (Sal) | 266.2 ± 4.4 |
| CQ | 249.8 ± 7.5 |
| Veh (DMSO) | 194.9 ± 20.9 |
| Baf | 193.3 ± 21.2 |
| Veh (Sal + DMSO) | 204.4 ± 8.4 |
| CQ+Baf | 281.7 ± 18.6* |
| Control (no veh) | 181.9 ± 3.7 |

Veh, vehicle; Sal, saline; CQ, chloroquine; Baf, bafilomycin. n=5-9/group. Related to Figure 2C. Data presented as mean ± SEM, \*p<0.05.

## Notes

### Competing Interest Statement

The authors have declared no competing interest.

